# Third dose COVID-19 mRNA vaccine enhances IgG4 isotype switching and recognition of Omicron subvariants by memory B cells after mRNA but not adenovirus priming

**DOI:** 10.1101/2023.09.15.557929

**Authors:** Gemma E. Hartley, Holly A. Fryer, Paul A. Gill, Irene Boo, Scott J. Bornheimer, P. Mark Hogarth, Heidi E. Drummer, Robyn E. O’Hehir, Emily S.J. Edwards, Menno C. van Zelm

## Abstract

**Background:** Booster vaccinations are recommended to improve protection against severe disease from SARS-CoV-2 infection. With primary vaccinations involving various adenoviral vector and mRNA-based formulations, it remains unclear if these differentially affect the immune response to booster doses. We here examined the effects of homologous (mRNA/mRNA) and heterologous (adenoviral vector/mRNA) vaccination on antibody and memory B cell (Bmem) responses against ancestral and Omicron subvariants.

**Methods:** Healthy adults who received primary BNT162b2 (mRNA) (n=18) or ChAdOx1 (vector) (n=25) vaccination were sampled 1-month and 6-months after their 2nd and 3rd dose (homologous or heterologous) vaccination. Recombinant spike receptor-binding domain (RBD) proteins from ancestral, Omicron BA.2 and BA.5 variants were produced for ELISA-based serology, and tetramerized for immunophenotyping of RBD-specific Bmem.

**Results:** Dose 3 boosters significantly increased ancestral RBD-specific plasma IgG and Bmem in both cohorts. Up to 80% of ancestral RBD-specific Bmem expressed IgG1^+^. IgG4^+^ Bmem were detectable after primary mRNA vaccination, and expanded significantly to 5-20% after dose 3, whereas heterologous boosting did not elicit IgG4^+^ Bmem. Recognition of Omicron BA.2 and BA.5 by ancestral RBD-specific plasma IgG increased from 20% to 60% after the 3rd dose in both cohorts. Reactivity of ancestral RBD-specific Bmem to Omicron BA.2 and BA.5 increased following a homologous booster from 40% to 60%, but not after a heterologous booster.

**Conclusion:** A 3rd mRNA dose generates similarly robust serological and Bmem responses in homologous and heterologous vaccination groups. The expansion of IgG4^+^ Bmem after mRNA priming might result from the unique vaccine formulation or dosing schedule affecting the Bmem response duration and antibody maturation.

## INTRODUCTION

Severe acute respiratory coronavirus-2 (SARS-CoV-2) causing the coronavirus disease-2019 (COVID-19) pandemic has resulted in over 750 million infections and over 6.9 million deaths (*1*). To combat the worldwide pandemic, the scientific community rapidly developed new vaccination technologies to reduce the burden of infections. Novel mRNA (BNT162b2 and mRNA-1273) and adenoviral vector (ChAdOx1 and Ad26.COV2.S) formulations were used in primary vaccination schedules across the globe (*2–6*) with high protection against severe disease (85-100%) (*6–8*). These vaccines generate responses to the SARS-CoV-2 spike protein, inducing antibodies directed to the receptor binding domain (RBD) that can prevent viral entry into host cells (*2–6*). Due to the nature of these vaccines to induce host cell expression of viral spike proteins, these elicit both high antibody titers and memory B cell (Bmem) numbers, as well as CD4^+^ and CD8^+^ T cell responses to protect against viral infection (*2, 4*).

In Australia, the initial primary vaccinations in 2021 were performed with two doses of BNT162b2 or ChAdOx1 with either a 3-4 week interval (*2*) or a 12-week interval (*5, 9*), respectively. Following the link between ChAdOx1 and vaccine-induced thrombocytopenia and thrombosis (*10, 11*), from April 2021 mRNA vaccinations (BNT162b2 or mRNA1273) were preferentially used in primary schedules and subsequent booster vaccinations. Due to border closures and COVID-19 restrictions, SARS-CoV-2 infection rates were low across Australia until late 2021 when the Delta subvariant caused a spike in infections. However, infection rates overall remained low, and the adult population had the opportunity to obtain 3 vaccine doses before more widespread infections with the Omicron variant (*12–14*).

Primary BNT162b2 vaccination generates robust antibody and Bmem responses with a predominant IgG1^+^ Bmem response (*14, 15*). The second dose also increased the reactivity of antibodies and Bmem to SARS-CoV-2 variants (*14, 15*). While the antibody response contracts after 1 month, Bmem numbers, their capacity to recognize viral variants, and their levels of somatic hypermutation (SHM) continue to increase up to 6-months post-vaccination (*16–18*). This suggests that there might be ongoing Bmem maturation due to continual germinal center (GC) activity (*16, 17*) which could be supported by spike-specific T follicular helper cells (Tfh), which remain stably present in GCs up to 6-months post-vaccination (*19*).

The immune response to primary adenoviral vector vaccination has not been as extensively studied. While it elicits significantly lower antibody levels than BNT162b2 vaccination (*12, 20*), it generates similar numbers of ancestral (Wuhan-Hu-1; WH1) RBD-specific Bmem numbers (*12*), which are also durable up to 6-months post-vaccination (*20, 21*). Still, to date there is little evidence to suggest that primary adenoviral vector vaccination induces continual Bmem maturation or GC activity.

Third dose booster vaccinations were successful in boosting protection against severe disease from SARS-CoV-2 variants including Omicron (*22–24*). In addition to higher serum IgG (*22, 25, 26*), a third dose mRNA booster was shown to increase the proportion of IgG-switched spike-specific Bmem (*22, 26*). Interestingly, this third dose booster also induced serum IgG4 and an expansion of Bmem expressing IgG4 (*26*). IgG4^+^ Bmem are mostly CD27^+^ and contain high levels of SHM, suggestive of an origin from secondary responses (*27*). As IgG4 responses are uncommon after other booster vaccinations (eg. Influenza) (*28*), it remains unclear whether this phenomenon is related to the antigen or to the vaccine formulation. Here, we addressed this by detailed evaluation of the antibody and Bmem response in individuals who received a homologous (primary mRNA with mRNA boost) or heterologous (primary adenoviral vector with mRNA boost) COVID-19 vaccination schedule.

## METHODS

### Participants

Healthy individuals without hematological or immunological disease, who had decided to take the COVID-19 vaccine were enrolled in a low-risk research study to examine their immune response to vaccination. Following informed consent, basic demographics (age and sex) were collected, as well as blood samples before and after each of three vaccinations between March 2021 and July 2022. The volunteers received either homologous (primary 2-dose BNT162b2 followed by BNT162b2 third dose, n = 18) or heterologous (primary 2-dose ChAdOx1 nCoV-19 followed by BNT162b2 third dose, n = 25) vaccinations. Of the 43 third dose boosters, one was mRNA-1273 and the other 42 were BNT162b2 (**Supplementary tables 1 and 2**). The cohorts were established previously, and responses were reported pre-vaccination, 3-4 weeks after dose 1 and 1 month after dose 2 (*12, 14*). For this study, samples were evaluated that were obtained 1 and 6 months after doses 2 and 3. This study was conducted according to the principles of the Declaration of Helsinki and approved by local human research ethics committees (Alfred Health ethics no. 32-21/Monash University project no. 72794).

### Sample processing

Blood samples were processed as described previously (*28–30*). Briefly, 200 µl was used for whole blood cell counts (Cell-Dyn analyzer; Abbott Core Laboratory, Abbott Park, IL) and Trucount analysis (see flow cytometry section). The remainder of the sample was used to separate and store plasma (-80°C), and to isolate live peripheral blood mononuclear cells (PBMC) following Ficoll-paque density gradient centrifugation and cryopreservation in liquid nitrogen for later analysis of RBD-specific B cells.

### Protein production and tetramerization

Recombinant spike RBD and nucleoprotein (NCP) proteins of the SARS-CoV-2 ancestral, Delta and Omicron BA.2 and BA.5 subvariant RBDs were produced with the N-terminal Fel d 1 leader sequence and C-terminal biotin ligase (BirA) AviTag and 6-His affinity tags, as described previously (*14, 30*). The RBD from the SARS-CoV-2 variants contained the following mutations: B.1.617.2 (Delta) L452R, T478K; B.1.1.529 (Omicron BA.2): G339D, S371F, S373P, S375F, S376A, D405N, R408S, K417N, N440K, S477N, T478K, E484A, Q493K, Q498R, N501Y, Y505H; B.1.1.529 (Omicron BA.5): BA.2 mutations with additional L452R, F486V and reversion of Q498. The DNA constructs were cloned into a pCR3 plasmid and produced and purified as described previously (*14, 30*). DNA was transfected into 293F cells using the Expi293 Expression system (Thermo Fisher Scientific, Waltham, MA). Following 5-day cultures at 37°C (ancestral and Delta) or 34°C (Omicron subvariants), harvested supernatants were purified using a Talon NTA-cobalt affinity column (Takara Bio, Kusatsu, Shiga, Japan) with elution in 200 mM Imidazole. Purified proteins were then dialyzed into 10 mM Tris and biotinylated (*14, 30*). Biotinylated proteins were subsequently dialyzed against 10 mM Tris for 36 hours at 4°C with 3 or more exchanges, and subsequently stored at -80°C prior to use. Soluble biotinylated RBD proteins were tetramerized with unique fluorochrome-conjugated streptavidins at a protein:streptavidin molar ratio of 4:1 to form: [RBD WH1]_4_-BUV395, [RBD WH1]_4_-BUV737, [RBD BA.2]_4_-BV480 and [RBD BA.5]_4_-BV650.

### Measurement of SARS-CoV-2 neutralizing antibodies in plasma

Measurement of neutralizing antibodies was performed using SARS-CoV-2 retroviral pseudotyped particles and a 293T-ACE2 cell line, as described previously (*14, 30*). Briefly, plasma was heat inactivated at 56°C for 45 minutes and serially diluted in DMF10. Diluted samples were then mixed with an equal volume of SARS-CoV-2 (Wuhan-1 Ancestral, Delta, BA.2 and BA.4/5 spike) retroviral pseudotyped virus and incubated for 1 hour at 37°C. Virus-plasma mixtures were added to 293T-ACE2 monolayers seeded the day prior at 10,000 cells/well, incubated for 2 hours at 37°C, before addition of an equal volume of DMF10 and incubated for 3 days. After incubation, tissue culture fluid was removed, and monolayers were washed once with PBS and lysed with cell culture lysis reagent (Promega, Madison, WI) and luciferase measured using luciferase substrate (Promega) in a Clariostar plate reader (BMG LabTechnologies, Offenburg, Germany). The percentage entry was calculated as described previously (*14, 30*), and plotted against reciprocal plasma dilution GraphPad Prism 9 Software (GraphPad Software, La Jolla, CA) and curves fitted with a one-site specific binding Hill plot. The reciprocal dilution of plasma required to prevent 50% virus entry was calculated from the non-linear regression line (ID50). The lowest amount of neutralizing antibody detectable is a titer of 20. All samples that did not reach 50% neutralization were assigned an arbitrary value of 10.

### ELISA

For quantification of total IgG against NCP and ancestral RBD and NCP, EIA/RIA plates (Costar, St Louis, MO) were coated with 2μg/ml recombinant SARS-CoV-2 ancestral RBD or NCP overnight at 4°C. Wells were blocked with 3% BSA in PBS and subsequently incubated with plasma samples. Plasma was diluted 1:300 for quantification of ancestral RBD- and NCP-specific antibodies post-dose 2, 6-months post-dose 2, post-dose 3 and 6-months post-dose 3. Plasma was titrated from 1:30 to 1:10,000 for quantification of ancestral and variant RBD-specific antibodies post-dose 2 and 3. Antigen-specific IgG was detected using rabbit anti-human IgG HRP (Dako, Glostrup, Denmark). ELISA plates were developed using TMB solution (Life Technologies, Carlsbad, CA) and the reaction was stopped with 1 M HCl. Absorbance (OD450nm) was measured using a Multiskan Microplate Spectrophotometer (Thermo Fisher Scientific). Serially diluted recombinant human IgG (in-house made human Rituximab) was used for quantification of specific IgG in separate wells on the same plate. Area under the curve (AUC) was calculated for each titration curve using GraphPad Prism software. Relative recognition of the RBD variants was calculated as a percentage of the AUC for that variant relative to the AUC for ancestral RBD.

For quantification of ancestral RBD-specific IgG1 and IgG4, EIA/RIA plates (Costar) were coated with 2 or 1 μg/ml recombinant SARS-CoV-2 ancestral RBD overnight at 4°C for IgG1 and IgG4 ELISAs respectively. Wells were blocked with 5% skim milk powder (SMP) in PBS and subsequently incubated with plasma samples. Plasma was diluted from 1:100 to 1:2000 for quantification of ancestral RBD-specific IgG1 and IgG4 antibodies post-dose 2, 6-months post-dose 2, post-dose 3 and 6-months post-dose 3 using mouse anti-human IgG1 biotin (Thermo Fisher Scientific) and mouse anti-human IgG4 biotin (Sigma-Aldrich, St Louis, MO), respectively. Finally, high sensitivity streptavidin HRP (Thermo Fisher Scientific) was added, and ELISA plates were developed using TMB solution (Life Technologies, Carlsbad, CA) and the reaction was stopped with 1 M HCl. Absorbance (OD450nm) was measured using a Multiskan Microplate Spectrophotometer (Thermo Fisher Scientific). Serially diluted recombinant human IgG1 or human IgG4 (BioRad, Hercules, CA) with unrelated specificities were used for quantification in separate wells on the same plate.

### Flow cytometry

Absolute numbers of leukocyte subsets were determined as previously described (*29, 30*). Briefly, 50 μl of whole blood was added to a Trucount tube (BD Biosciences) together with 20 μl of antibody cocktail containing antibodies to CD3, CD4, CD8, CD16, CD19, CD56 and CD45 from the 6-color TBNK reagent kit (BD Biosciences) (**Supplementary Tables 4 and 5**) and incubated for 15 minutes at room temperature in the dark. Subsequently, samples were incubated for a further 15 minutes at room temperature with 500 μl of 1X BD Lysis solution (BD Biosciences) to lyse red blood cells. The tube was then stored in the dark at 4^°^C for up to 2 hours prior to acquisition on a LSRII or FACSLyric analyzer (BD Biosciences).

For the detection of antigen-specific Bmem, cryopreserved PBMC were thawed and stained as previously described (*12, 14, 30*). Briefly, 10-15 million PBMC were incubated with fixable viability stain 700 (BD Biosciences), antibodies against CD3, CD19, CD21, CD27, CD38, CD71, IgA, IgD, IgG1, IgG2, IgG3, IgG4, (**Supplementary Tables 4 and 5**) and 5 μg/ml each of [RBD WH1]_4_-BUV395, [RBD WH1]_4_-BUV737, and [RBD BA.2]_4_-BV480 and [RBD BA.5]_4_-BV650 for 15 minutes at room temperature in a total volume of 250 μl FACS buffer (0.1% sodium azide, 0.2% BSA in PBS). In addition, 5 million PBMC were similarly incubated with fixable viability stain 700 (BD Biosciences), antibodies against CD3, CD19, CD27 and IgD, and BUV395-, BUV737-, BV480- and BV650-conjugated streptavidin controls (**Supplementary Tables 4 and 5**). Following staining, cells were washed with FACS buffer, fixed with 2% paraformaldehyde for 20 minutes at room temperature and washed once more. Following filtration through a 70 μM filter, cells were acquired on a 5-laser LSRFortessa X-20 (BD Biosciences). Flow cytometer set-up and calibration was performed using standardized EuroFlow SOPs, as previously described (**Supplementary Tables 6 and 7**) (*31*).

### Data analysis and statistics

All flow cytometry data were analyzed with FlowJo v10 software (BD Biosciences). Statistical analysis was performed with GraphPad Prism 9 Software (GraphPad Software). Matched pairs were analyzed with the non-parametric Wilcoxon matched pairs signed rank test with Bonferroni correction for multiple comparisons. Comparisons between 3 or more groups were performed using the Friedman’s test (paired) or Kruskal-Wallis (unpaired) with Dunn multiple comparisons test. For all tests, *p* < 0.05 was considered significant.

## RESULTS

### Third dose booster increases ancestral RBD-specific Bmem irrespective of primary vaccination formulation

Blood samples were collected at 1 and 6 months after both dose 2 (D2 and 6mD2, respectively) and dose 3 (D3 and 6mD3, respectively) from 18 individuals who received a homologous vaccination schedule (3x mRNA) and 25 individuals who received heterologous vaccination (2x ChAdOx1, 1x mRNA) (**Figure 1A, Supplementary Tables 1, 2, 3**). There were no significant differences in sampling times between the two cohorts apart from 6-months post-dose 2: 185 (homologous) vs 178 days (heterologous) (*p* < 0.0001) (**Supplementary Table 3**). The cohorts did not differ in age, but the homologous group trended to include fewer females (56%) than the heterologous group (80%; *p* = 0.09) (**Supplementary Tables 1 and 2**).

**Figure 1:**
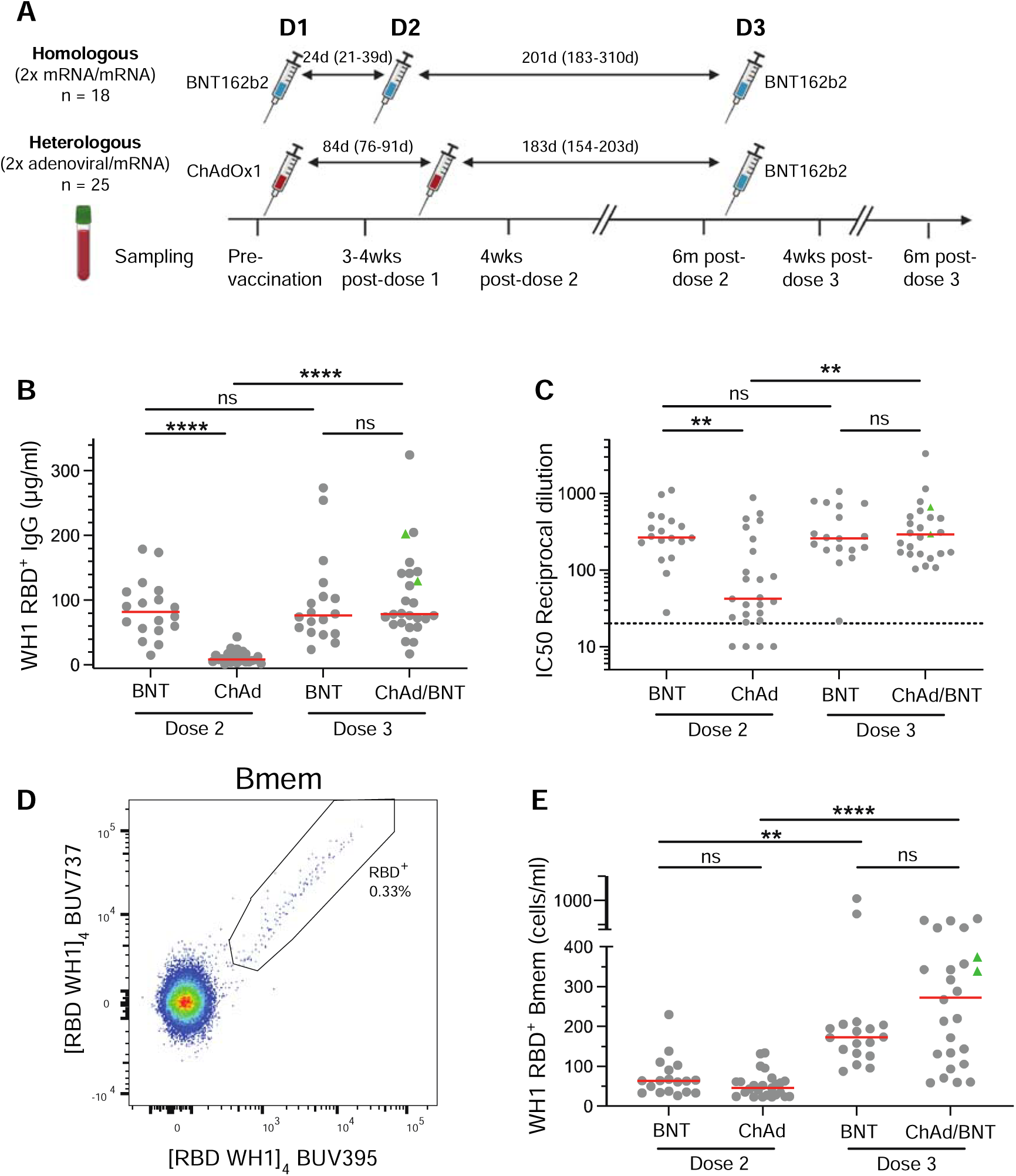
Third dose booster significantly increases WH1 RBD-specific plasma IgG and Bmem. (**A**) Schematic of patient cohorts, vaccination schedules and sampling timepoints. Details in **Supplementary Tables 1-3**. (**B**) Ancestral (WH1) RBD-specific plasma IgG levels and (**C**) neutralizing antibodies (NAb) in individuals who received a primary BNT162b2 (BNT) or ChAdOx1 (ChAd) vaccination followed by an mRNA third dose booster. (**D**) Detection of ancestral RBD-specific Bmem using double discrimination with recombinant WH1 RBD tetramers. (**E**) WH1 RBD-specific Bmem numbers following 2 doses of BNT or ChAd and after mRNA third dose booster. Green triangles represent individuals who had a confirmed breakthrough infection (BTI) prior to sampling (**Supplementary Tables 1 and 2**). Red lines in panels B, C and E represent median values. Kruskal-Wallis test with Dunn’s multiple comparisons test. ** p < 0.01, **** p < 0.0001.

The vaccine-specific antibody and Bmem responses were evaluated using recombinantly-produced RBD proteins of ancestral and Omicron BA.2 and BA.5, whereas SARS-CoV-2 nucleocapsid protein (NCP)-specific plasma IgG and was evaluated to confirm self-reported breakthrough infections (BTI) (*12, 14, 30*). As previously reported (*12, 14*), our BNT162b2 cohort had 8-10-fold higher ancestral RBD-specific plasma IgG and neutralizing antibodies (NAb) than the ChAdOx1 cohort at 1-month post-dose 2 (**Figure 1B, C**). The third dose mRNA booster elicited similar RBD-specific IgG and NAb responses in both cohorts, with levels comparable to those of the mRNA cohort after dose 2 (**Figure 1B, C**). Importantly, all donors generated detectable NAb after dose 3, even those four that did not reach neutralizing capacity after 2 doses of ChAdOx1 (**Figure 1C**).

Ancestral RBD-specific Bmem were evaluated within CD19^+^ B cells after exclusion of CD27^-^IgD^+^ naive B cells (**Supplementary** Figure 1) through double discrimination, i.e. positivity for both [RBD WH1]_4_-BUV395 and [RBD WH1]_4_-BUV737 (**Figure 1D**). The primary BNT162b2 and ChAdOx1 vaccinated cohorts had similar ancestral RBD-specific Bmem numbers after dose 2 (**Figure 1E**) (*12*). The third mRNA dose significantly increased ancestral RBD-specific Bmem numbers in both cohorts irrespective of the primary schedule (**Figure 1E**).

### Durability of ancestral RBD-specific Bmem up to 6 months after 2 and 3 vaccine doses

To evaluate the durability of the response, the vaccine-induced antibody levels and Bmem numbers were quantified and compared between 1- and 6-months post-dose 2 and 3. Multiple participants self-reported SARS-CoV-2 BTI, and these were confirmed with NCP-specific plasma IgG (*12, 14*) (**Supplementary Tables 1 and 2**) (**Figure 2 A, E**). These samples are marked (green triangles) to visualize a potential confounding effect (**Figure 2A-H**). In line with previous observations, the plasma IgG and NAb levels contracted between 1 and 6 months after both dose 2 and dose 3 (**Figure 2B,C,F,G**). Within the complete cohorts, the contractions were not significant after dose 3. This was due to a confounding effect of BTIs: Following stratification, this decline was significant for the groups without BTI (**Figure 2B, F**).

**Figure 2:**
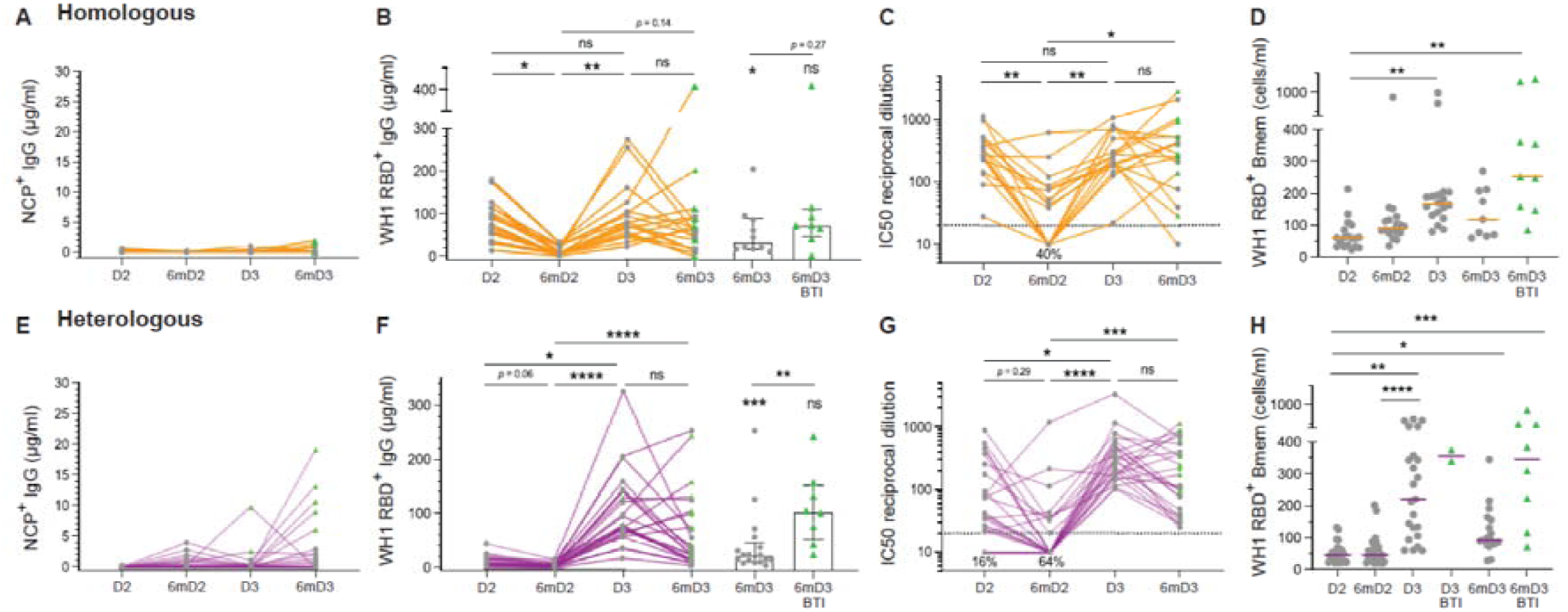
Ancestral (WH1) RBD-specific plasma IgG and Bmem dynamics following homologous or heterologous vaccination. (**A**) NCP-specific, (**B**) WH1 RBD-specific, (**C**) Neutralizing antibody (NAb) levels and (**D**) WH1 RBD-specific Bmem numbers following homologous vaccination (n = 18). (**E**) NCP-specific, (**F**) WH1 RBD-specific, (**G**) NAb levels and (**H**) WH1 RBD-specific Bmem numbers following heterologous vaccination (n = 25). Green triangles represent individuals who had a confirmed breakthrough infection (BTI) prior to sampling (**Supplementary Tables 1 and 2**). Red lines in panels D and H represent median values. Significance stars above bars in B and F depict comparisons to the 1-month post-dose 3 measurements. Kruskal-Wallis test with Dunn’s multiple comparisons test. * *p* < 0.05, ** *p <* 0.01, **** *p <* 0.001, **** *p <* 0.0001

Double dose primary BNT162b2 vaccination generated a population of ancestral RBD-specific Bmem that trended to increase in number at 6-months post-dose 2 (prior to the third dose), however, this was not seen in the heterologous cohort (**Figure 2D, H)**. The third dose significantly boosted ancestral RBD-specific Bmem numbers in both cohorts with heterologous vaccination generating a wider range of Bmem numbers (**Figure 2D, H**). Individuals who had BTIs between 1- and 6-months post-dose 3 trended to have higher numbers of ancestral RBD-specific Bmem at 6-months post-dose 3. No significant decline of ancestral RBD-specific Bmem numbers was observed in SARS-CoV-2 naive individuals at 6-months post-dose 3 in either cohort (**Figure 2D, H**). In summary, irrespective of primary vaccination, the antibody responses contract between 1 and 6 months after doses 2 and 3, whereas the ancestral RBD-specific Bmem numbers remain more stable.

### Transient expansion of recently activated ancestral RBD-specific Bmem at 1 month after mRNA vaccination

To evaluate the maturation status of ancestral RBD-specific Bmem, these were further evaluated for surface marker expression (**Figure 3A**). CD71 is expressed on recently activated cells to provide uptake of iron for proliferation (*32, 33*) and is typically downregulated within 14 days (*30, 34*). Both cohorts at all timepoints had only minor fractions and numbers of vaccine induced RBD-specific Bmem (median <3%) expressing CD71, indicative of quiescent populations (**Figure 3B, Supplementary** Figure 2A). The exceptions were 3 samples in the heterologous cohort that were obtained within 15 days of BTI (participants no. 20, 34 and 42) (**Figure 3B, Supplementary Table 2**). Low expression of CD21 on Bmem is another marker of recent activation (*35*). While the majority of RBD-specific Bmem at 1 month after doses 2 and 3 were CD21^+^, frequencies of CD21^lo^ cells significantly increased at 1 month and then significantly declined at 6-months after each dose (**Figure 3C, Supplementary** Figure 2B). Finally, expression of CD27 was evaluated within RBD-specific IgG^+^ Bmem as a marker for more mature IgG^+^ Bmem (*36, 37*). The majority of RBD-specific IgG^+^ Bmem were CD27^+^ 1 month after dose 2 with was no significant difference in frequency between cohorts at this timepoint (*p* = 0.23). However, frequencies significantly increased 6 months after dose 2 and 3 mRNA vaccination, but not 6 months after dose 2 adenoviral vector vaccination (**Figure 3D**). Together this phenotypic evaluation demonstrates that at 1 month after each vaccine, the RBD-specific Bmem populations do not display signs of recent proliferation, similar to the total Bmem compartment (**Supplementary** Figure 3). Still, the RBD-specific Bmem at 1-month post-dose 2 and dose 3 contain large fractions of recently activated cells that further mature by 6 months, thereby gaining CD21 and CD27 expression. These signs of further maturation between 1 and 6 months are particularly notable following mRNA vaccination.

**Figure 3:**
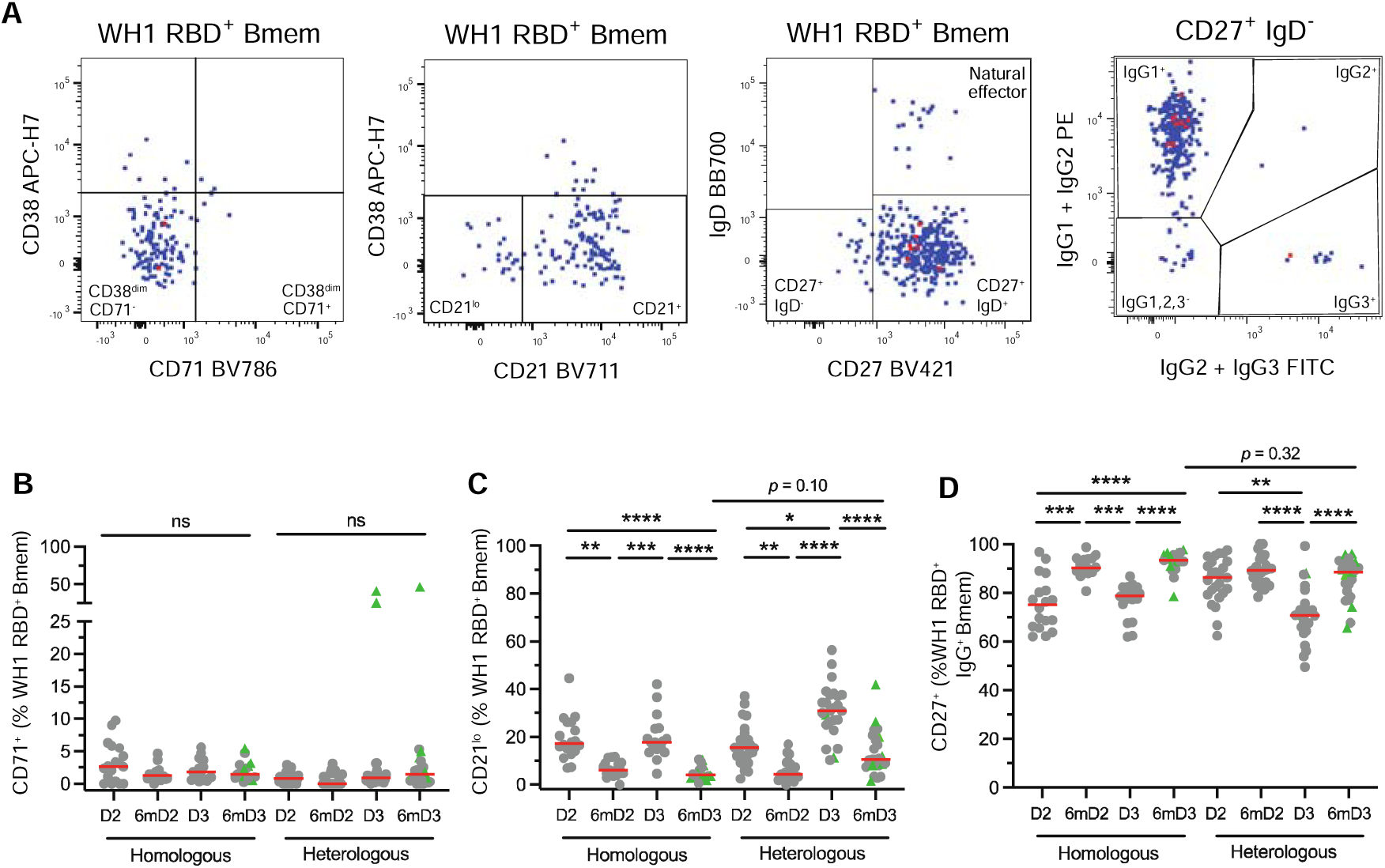
mRNA third dose generates a transient population of recently activated ancestral (WH1) RBD-specific Bmem. (**A**) Definitions of CD38^dim^CD71^+^, CD21^lo^ and CD27^+^IgG^+^ WH1 RBD-specific Bmem. Proportions of WH1 RBD-specific (**B**) CD38^dim^CD71^+^ Bmem (**C**) CD21^lo^ Bmem (**D**) and CD27^+^IgG^+^ Bmem in the homologous (n = 18) and heterologous (n = 25) vaccination groups. Green triangles represent individuals who had a confirmed breakthrough infection (BTI) prior to sampling (**Supplementary Tables 1 and 2**). Red lines in panels B-D represent median values. Kruskal-Wallis test with Dunn’s multiple comparisons test. * *p* < 0.05, ** *p* < 0.01, *** *p* < 0.001, **** *p* < 0.0001.

### Expansion of IgG4^+^ Bmem after homologous boost is absent from heterologous boost

To investigate whether the previously observed IgG4 response (*26, 38*) also occurred in our cohorts, we evaluated the plasma IgG1 and IgG4 subclass contributions to the homologous or heterologous third dose booster responses. Both cohorts generated high levels of ancestral RBD-specific plasma IgG1 after the third dose, and these recapitulated the dynamics of the total ancestral RBD-specific IgG (**Figure 4A**). Ancestral RBD-specific plasma IgG4 was detectable at low level in the mRNA primed cohort, and these levels were boosted after the third dose (**Figure 4B**). In contrast, the adenoviral vector primed cohort showed very little IgG4 prior to and after the third dose boost (**Figure 4B**). Thus, priming with an mRNA or adenoviral vector vaccine has differential effects on the capacity of recipients to form IgG4 responses.

**Figure 4:**
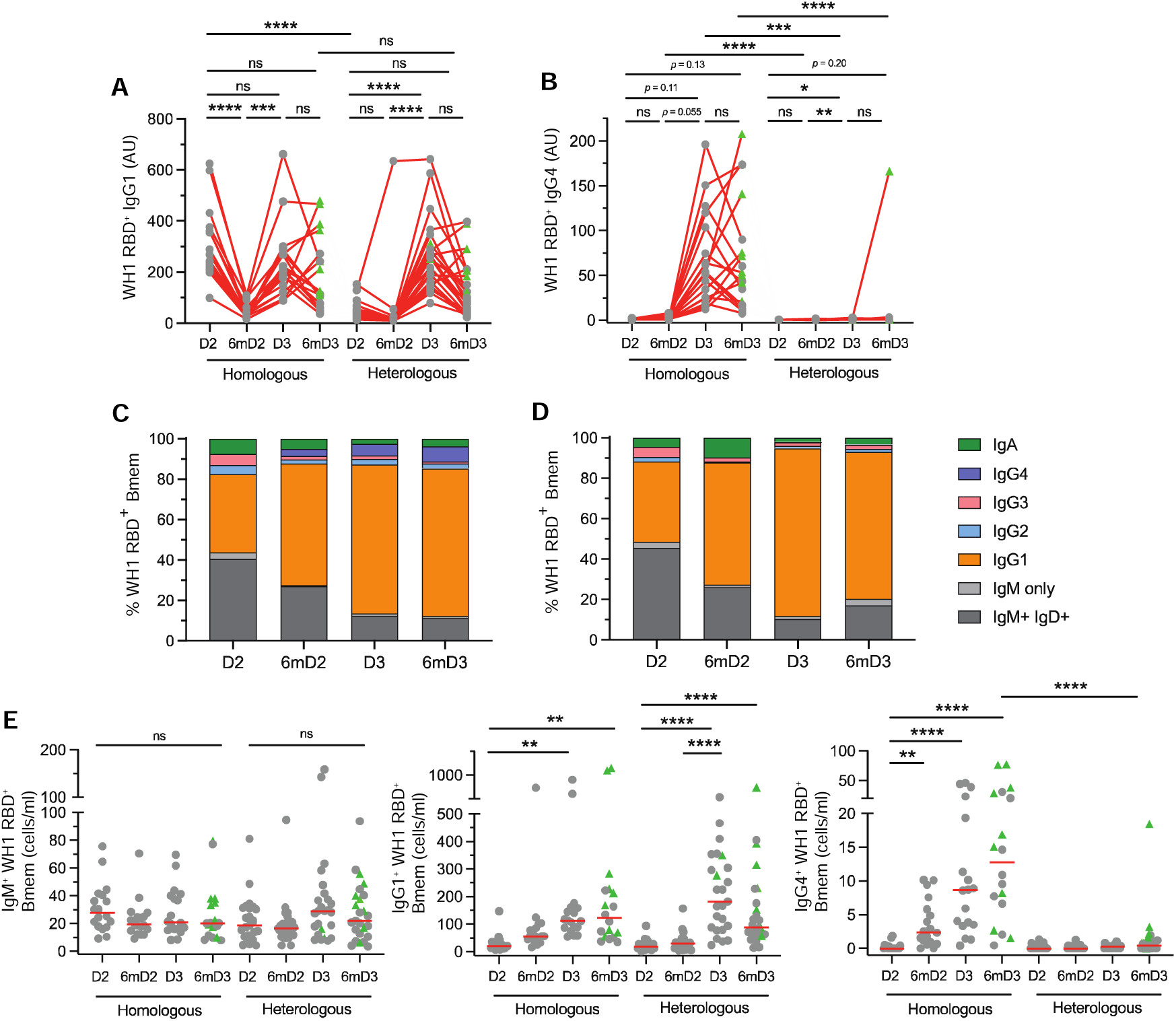
Predominant IgG1 response with IgG4^+^ Bmem population enhanced in mRNA-primed cohort. Levels of ancestral (WH1) RBD-specific plasma (**A**) IgG1 and (**B**) IgG4. Median frequencies of WH1 RBD-specific Bmem in the (**C**) homologous or (**D**) heterologous vaccination groups. (**E**) IgM^+^, IgG1^+^ and IgG4^+^ WH1 RBD-specific Bmem numbers. Green triangles represent individuals who had a confirmed breakthrough infection (BTI) prior to sampling (**Supplementary Tables 1 and 2**). Red lines in panel C represent median values. Kruskal-Wallis test with Dunn’s multiple comparisons test. * *p* < 0.05, ** *p* < 0.01, *** *p* < 0.001, **** *p* < 0.0001

IgG4^+^ Bmem are presumed to be formed after ongoing GC responses or after renewed encounters with the same antigen (*27*). Therefore, we here evaluated vaccine-elicited formation of IgG4^+^ Bmem in the two cohorts before and after the third dose booster. Through our extensive immunophenotyping including Ig isotypes and IgG subclasses, we previously found expansions of IgG1^+^, but not IgG4^+^ Bmem after 2 doses of BNT162b2 or ChAdOx1 in COVID-19 naïve individuals (*12, 14*). With the same approach (**Supplementary** Figure 1A**, B**), we here found that in both cohorts the ancestral RBD-specific IgG1^+^ Bmem population remained stable at 6-months post-dose 2 and was further expanded after dose 3 (**Figure 4C-E, Supplementary** Figures 1C**, 4 and 5**). The IgM^+^ Bmem and IgA^+^ Bmem populations were not affected (**Fig 4C-E, Supplementary** Figures 4-6). At 6-months post-dose 3, the IgG1 compartment encompassed approximately 80-90% of the ancestral RBD-specific Bmem compartment (**Figure 4C, D**). This enrichment was not apparent in the total Bmem compartment, in which only about 20% expressed IgG1 (**Supplementary Figure 7**). Importantly, the BNT162b2-primed cohort showed a significant enlargement of the ancestral RBD-specific IgG4^+^ Bmem compartment at 6-months post-dose 2 (**Figure 4C, E, Supplementary Figure 1D**). This population was further expanded at 1- and 6-months post-dose 3. In contrast, the ChAdOx1-primed cohort had very few ancestral RBD-specific IgG4^+^ Bmem at each timepoint (**Figure 4D, E**), except for one individual who had a BTI prior to the 6-months post-dose 3 sample. Thus, the expansion of plasma IgG4 and IgG4^+^ Bmem after a third dose booster is restricted to the mRNA-primed cohort, suggesting that either the primary vaccination formulation or the unique primary dosing schedule (3-week interval) underlies this.

### An mRNA booster enhances recognition of Omicron subvariants by Bmem irrespective of primary vaccination

To examine the effect of mRNA booster vaccination on recognition of SARS-CoV-2 variants, we evaluated the capacity of plasma antibodies and ancestral-RBD-specific Bmem to recognize Delta (B.1.617.2) and Omicron (B.1.1.529) BA.2 and BA.5 variant RBD proteins. NAb levels to all three variants were significantly higher at 1-month post-dose 3 than at 1-month post-dose 2 for both cohorts (**Figure 5A, B**). Importantly, after dose 3, plasma from all donors had the capacity to neutralize all variants, whereas only all individuals generated NAbs to Delta after 2 doses of BNT162b2(**Figure 5A, B**). Plasma RBD-binding serology was performed to evaluate the relative capacity of ancestral RBD-specific IgG to bind each variant. In line with previous findings, the capacity of ancestral RBD-specific plasma IgG to bind Delta RBD was almost 100% (**Figure 5C, D**). In contrast, the median recognition of BA.2 and BA.5 was <30% after 2 vaccine doses, irrespective of formulation, and these increased significantly after the third dose mRNA booster to 50-60% (**Figure 5C, D**).

**Figure 5:**
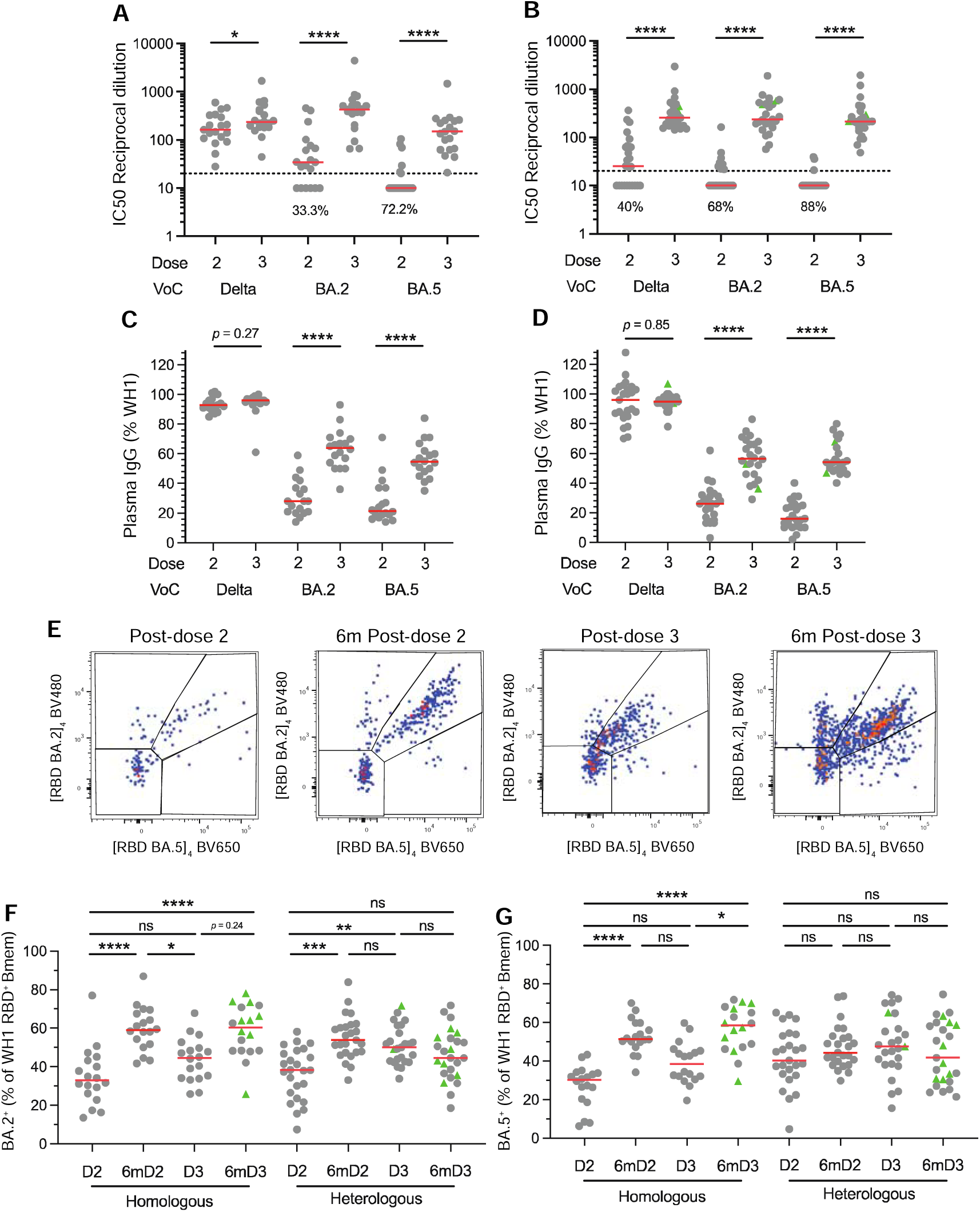
Third dose booster vaccination increases recognition of Omicron BA.2 and BA.5 variants. Neutralization of Delta and Omicron BA.2 and BA.5 variants in the (**A**) homologous and (**B**) heterologous vaccination groups. Recognition of ancestral (WH1) RBD-specific plasma IgG to Delta and Omicron BA.2 and BA.5 variants in the (**C**) homologous and (**D**) heterologous vaccination groups. (**E**) Identification of WH1 RBD-specific Bmem that also bind Omicron BA.2 and BA.5 variants. Capacity of WH1 RBD-specific Bmem to recognise Omicron (**F**) BA.2 and (**G**) BA.5 in the following homologous (n = 18) and heterologous vaccination groups (n = 25). Green triangles represent individuals who had a confirmed breakthrough infection (BTI) prior to sampling (**Supplementary Tables 1 and 2**). Red lines in panels C and D represent median values. Wilcoxon signed rank test with Bonferroni correction for multiple comparisons for panels A-D. Kruskal-Wallis test with Dunn’s multiple comparisons test for F and G. * *p* < 0.05, ** *p* < 0.01, **** *p* < 0.0001.

In addition, we used fluorescent tetramers of Omicron BA.2 and BA.5 RBDs to evaluate the capacity of ancestral RBD-specific Bmem to bind either or both subvariants (**Figure 5E**). Around 30-40% of ancestral RBD-specific Bmem bound BA.2 at 1-month post-dose 2 in both cohorts, and this recognition significantly increased to 60% at 6-months post-dose 2 (**Figure 5F**). BA.2 recognition 1-month post-dose 3 was about 50% and this increased in the homologous cohort to 60% at 6-months post-dose 3, whereas no change was observed at 6-months post-dose 3 in the heterologous cohort (**Figure 5F**). Recognition of BA.5 showed similar patterns as BA.2 in the homologous cohort with significant increases from 30% to 50% at 1-month and 6-months post-dose 2, as well as from 40% to 60% at 1- and 6-months post-dose 3 (**Figure 5G**). In contrast, no significant changes were found for recognition of BA.5 in the heterologous group with median recognition in the range of 40-50% (**Figure 5G**). Variant-binding ancestral RBD-specific Bmem also showed a similar phenotype (ie. predominantly IgG1^+^ with IgG enrichment in the homologous group) to total ancestral RBD-specific Bmem in each respective vaccination cohort (**Supplementary** Figure 8).

In summary, we confirm previous findings that a third dose mRNA booster significantly expands ancestral RBD-specific plasma IgG and Bmem levels, irrespective of the primary vaccination schedule. Importantly, we expanded on previous observations that the formation of plasma IgG4 and IgG4^+^ Bmem is restricted to mRNA-primed individuals, and not present in adenoviral vector vaccine-primed individuals who are COVID-19 naive. While Bmem at 1-month post-vaccination appear to be quiescent, these do show signs of recent activation which are absent at 6-months post-vaccination. This ongoing maturation is associated with increased recognition of Omicron variants and is especially apparent after mRNA vaccination.

## DISCUSSION

We have shown that both homologous and heterologous COVID-19 booster vaccinations significantly increase ancestral RBD-specific plasma IgG and Bmem. An mRNA third dose induces a population of recently activated Bmem, but these contract at 6-months post-dose 3. The ancestral RBD-specific Bmem population seemed to further mature with an increase in the proportion of IgG^+^ Bmem that expressed CD27 at 6-months post-dose 3. Only after mRNA priming, a population of IgG4^+^ RBD-specific Bmem was apparent that expanded after the homologous third dose boost with increased recognition of Omicron BA.2 and BA.5 subvariants.

A third COVID-19 vaccine dose was recommended in late 2021 to all vaccinees after evidence suggested that antibody levels declined beyond 6-months post-primary vaccination and the risk of BTI increased (*39–42*). We confirm previously reported findings that a third dose (either homologous or heterologous) significantly boosts RBD-specific IgG and NAb levels regardless of primary vaccination formulation (*22, 43–45*). Ancestral RBD-specific Bmem numbers are also significantly increased following the third dose. We have previously shown that primary vaccination with either BNT162b2 or ChAdOx1 generated similar numbers of ancestral RBD-specific Bmem (*12*). We have expanded on this by demonstrating that in contrast to ancestral RBD-specific plasma IgG and NAb levels, ancestral RBD-specific Bmem numbers did not significantly decline at 6-months post-dose 3, in line with other studies (*16, 22, 46, 47*).

Previous studies have used increased CD71 expression and reduced CD21 expression as markers of recently activated Bmem (*34, 35*). The observed dynamics of CD71 expression in this study are in line with our previous findings following primary COVID-19 vaccination and others post-influenza vaccination, with CD71^+^ Bmem contracting beyond 7 days post-antigen exposure (*12, 14, 34*). Heterologous vaccination induces a significant expansion of CD21^lo^ ancestral RBD-specific Bmem that is still apparent 4 weeks after the third dose (30-40% of the compartment). In previous studies, we have shown that less than 30% of ancestral RBD-specific Bmem were CD21^lo^ 4 weeks after primary (double-dose) COVID-19 vaccination (*12, 14*). However, others have shown that this CD21^lo^ activated memory population can make up to 40-50% of the antigen-specific Bmem population 4 weeks after either influenza or COVID-19 vaccination, or SARS-CoV-2 infection, and does not contract to below 25% until after 90 days following vaccination (*35, 48*). Therefore, although the frequencies of CD21^lo^ Bmem observed following a third dose are higher than we have observed following primary COVID-19 vaccination, these levels are consistent with previous studies from other groups (*34, 35*).

Following the third dose booster, the frequencies of IgG^+^ Bmem that expressed CD27 initially declined at 1-month, followed by a significant increase at 6 months. Thus, the initial vaccine elicited Bmem population continued to mature for several months after mRNA booster vaccination. As CD27^+^IgG^+^ Bmem have higher levels of SHM and an increased replication history than CD27^-^IgG^+^ Bmem, these likely originate from ongoing or renewed GC reactions (*36, 37*). The CD21^lo^ Bmem compartment can comprise of cells either primed for plasma cell differentiation (CD21^lo^CD27^+^) (*49*) or from extrafollicular responses (CD21^lo^CD27^-^) (*48, 50*), and are thus unlikely to contribute to the long-term stability of the Bmem compartment. Hence, we infer that the increase in the proportion of CD27^+^IgG^+^ Bmem is indicative of continual GC activity and maturation beyond 1-month post-mRNA booster vaccination. Continual GC responses can generate a high affinity resting Bmem pool, which is important to maintain durable protection while recently activated Bmem numbers continue to decrease (*51, 52*).

Primary COVID-19 vaccination generated a predominant IgG1^+^ population of ancestral RBD-specific Bmem in both cohorts, which was further expanded after a third dose boost. Importantly, the expansion of IgG1^+^ Bmem was not at the expense of IgM^+^ Bmem, which remained present in similar numbers. IgG1^+^ Bmem can provide protection against BTI by secreting IgG1 antibodies upon re-exposure. IgG1 antibodies are potent neutralizers and are effective at activating the classical complement pathway and engaging Fc-mediated responses such as antibody-dependent cellular cytotoxicity and hence are important in the clearance of viral infections (*53, 54*). Thus, a predominant IgG1 response following COVID-19 vaccination is suitable for neutralization of this pathogen.

We here found that the significant expansion of IgG4^+^ Bmem after the third dose boost (*26*) was only apparent in the mRNA-primed cohort and not in the adenoviral vector-primed group. Thus, it raises the question as to the mechanisms that drive the formation of IgG4-secreting plasma cells and IgG4^+^ Bmem. Other vaccinations have generated the production of plasma IgG4 such as VAX003 (HIV), EBA-175 (Malaria) and acellular pertussis vaccinations (*55–57*). It is not well understood why these vaccine formulations and schedules induce IgG4 antibodies. However, it is worth noting that VAX003 has a multiple-dose schedule (7 doses) in comparison to other HIV vaccine candidates that are given as a single dose (*55*). In addition, IgG4 antibodies are also produced in response to the repeated antigen exposure of allergen immunotherapy (*58, 59*). Given the unique capacity of IgG4 antibodies to undergo Fab arm exchange, there could be a unique functional effect elicited by mRNA vaccination, although it should be noted that IgG1 still predominates the response.

Primary BNT162b2 vaccination has a shorter window between dose 1 and dose 2 (3 weeks) (*2*) compared to ChAdOx1 (12 weeks) (*5, 9*). As both groups received an mRNA third dose 6-months post-primary vaccination, the IgG4 expansion in the homologous group is either due to the timing between dose 1 and 2 or the primary vaccination formulation. Examining a cohort of individuals who received a primary mRNA vaccination with a longer duration between doses 1 and 2 would confirm whether this effect is due to dosing or vaccination formulation. In IgG4-related disease and Kimura disease, prominent IgG4 class switching is thought to be controlled by a population of Tfh cells co-expressing CXCR5, PD-1, ICOSL, IL-10, IL-4 and LAG-3 (*60, 61*). It would be of interest whether such a Tfh cell population is specifically generated by mRNA vaccination and/or repeat vaccinations with a short time interval (<1 month).

Whilst ancestral RBD-specific plasma IgG levels were increased at 1 month following the third vaccine dose, these levels had significantly declined at 6 months in individuals without a confirmed BTI. Individuals with a confirmed BTI at that timepoint showed an increase in RBD-specific plasma IgG and trended to have more ancestral RBD-specific Bmem. Others have also demonstrated that BTI following COVID-19 vaccination generates higher antibody and Bmem responses to COVID-19 naive vaccinated individuals (*47, 62, 63*). This suggests that subsequent antigen exposures, either through vaccination or infection, would continue to increase antibody and Bmem levels. However, we currently do not know if there are certain levels of antibodies or B and T cell numbers required to confer protection. What may be more important is not the overall boosting of the response, but enhanced recognition of SARS-CoV-2 variants, that may prevent infection and/or severe disease.

Homologous vaccination not only induces an IgG4 expansion, but also significantly increases recognition of RBD-specific Bmem to Omicron BA.2 and BA.5 up to 6-months post-dose 3, whereas heterologous vaccination induces limited improvement in recognition of Omicron subvariants. In contrast, both vaccination schedules showed an increase in NAb and RBD-specific plasma IgG recognition of Omicron BA.2 and BA.5. This suggests that upon receipt of the third dose, a number of pre-existing Bmem with a higher affinity to Omicron BA.2 and BA.5 may have differentiated into antibody-secreting cells and hence increased circulating antibody recognition of Omicron subvariants (*64*). Pre-existing Bmem can not only differentiate into plasmablasts but also re-enter the GC where they undergo further SHM and increase in affinity (*64*). mRNA primary vaccination induces continual GC activity resulting in an increase in Bmem and SHM levels (*16, 43, 65*). Therefore, the increase in SHM of ancestral RBD-specific Bmem elicited by homologous vaccination may increase the affinity of the B cell receptor, allowing some variant RBD mutations impacting binding to be overcome; however, further molecular studies are required to confirm this.

In summary, we have shown that the Bmem response elicited by a third dose booster with an mRNA vaccine is differentially affected by the primary vaccination (schedule and/or formulation). Both homologous and heterologous vaccine boosters significantly increased ancestral RBD-specific plasma IgG, NAbs, and Bmem numbers to a similar degree. Through extensive immunophenotyping, we show that ancestral RBD-specific Bmem show signatures of continual maturation for at least 6-months post-dose 3. However, homologous mRNA vaccination alone induces an expansion of ancestral RBD-specific IgG4-switched Bmem and an increased recognition of Omicron BA.2 and BA.5 by ancestral RBD-specific Bmem. It is still unclear whether IgG4 is having a supportive or inhibitory role in responses to subsequent boosters and what role this isotype plays in protection against disease. mRNA and adenoviral vector vaccines have only been widely utilized for the first time to combat the SARS-CoV-2 pandemic. Their rapid production rates and high efficacies make these ideal formulations to use against future pathogens. Our studies reveal how antibody and Bmem responses are generated to each vaccination type as well as booster doses and reveal important differences generated by each vaccine type. These vaccine technologies may be adopted to combat other pathogens in the future and these data provide further crucial evidence to help public health officials make informed recommendations about vaccination schedules and booter doses in the future.

## ACKNOWLEDGEMENTS

We thank Dr. Bruce D. Wines and Ms. Sandra Esparon (Burnet Institute) for technical assistance, Ms. Shir Sun, Mr. Jack Edwards and Ms. Ebony Blight (Monash University) for sample collection and preparation, and the staff of ARAFlowcore for flow cytometry support. Supported by an Australian Government Research Training Program Scholarship (GEH), the Australian Government Medical Research Future Fund (MRFF, Project no. 2016108; MCvZ, HED and REO’H) and an unrestricted research grant from BD Biosciences.

## AUTHOR CONTRIBUTIONS

Designed and/or performed experiments: GEH, HAF, PAG, IB, SJB, PMH, HED, REO’H, ESJE and MCvZ; Formal analysis: GEH, HAF, IB; Provided reagents: SJB; Supervised the work: ESJE, REO’H and MCvZ; Wrote the manuscript: GEH and MCvZ. All authors edited and approved the final version of the manuscript.

