## Supplementary Table 1 for "Third dose COVID-19 mRNA vaccine enhances IgG4 isotype switching and recognition of Omicron subvariants by memory B cells after mRNA but not adenovirus priming"

**Supplementary Tables** (n= 7) **and Figures** (n= 8)

**Table 1. Participants who received a homologous vaccination schedule**

| **Participant** | **Age (yrs)** | **Sex** | **Primary vaccine** | **Third dose vaccine** | **Timing of confirmed SARS-CoV-2 infection** |
| --- | --- | --- | --- | --- | --- |
| 1 | 24 | F | BNT162b2 | BNT162b2 | 81 days before 6-months post-dose 3 sample |
| 2 | 25 | F | BNT162b2 | BNT162b2 | 30 days before 6-months post-dose 3 sample |
| 3 | 25 | M | BNT162b2 | BNT162b2 | NA |
| 4 | 26 | M | BNT162b2 | BNT162b2 | NA |
| 5 | 28 | F | BNT162b2 | BNT162b2 | 29 days before 6-months post-dose 3 sample |
| 6 | 31 | M | BNT162b2 | BNT162b2 | NA |
| 7 | 32 | F | BNT162b2 | BNT162b2 | 66 days before 6-months post-dose 3 sample |
| 8 | 35 | F | BNT162b2 | BNT162b2 | NA |
| 9 | 37 | F | BNT162b2 | BNT162b2 | Between 1- and 6-months post-dose 3 |
| 10 | 38 | F | BNT162b2 | BNT162b2 | NA |
| 11 | 39 | M | BNT162b2 | BNT162b2 | 20 days before 6-months post-dose 3 sample |
| 12 | 40 | M | BNT162b2 | BNT162b2 | NA |
| 13 | 42 | F | BNT162b2 | BNT162b2 | 15 days before 6-months post-dose 3 sample |
| 14 | 45 | F | BNT162b2 | BNT162b2 | NA |
| 15 | 46 | M | BNT162b2 | BNT162b2 | NA |
| 16 | 48 | M | BNT162b2 | BNT162b2 | 119 days before 6-months post-dose 3 sample |
| 17 | 51 | F | BNT162b2 | BNT162b2 | 119 days before 6-months post-dose 3 sample |
| 18 | 62 | M | BNT162b2 | BNT162b2 | NA |
|  | 37 (24-62)  median w range | 56% Female |  | 100% BNT162b2 |  |

**Table 2. Participants who received a heterologous vaccination schedule**

| **Participant** | **Age** | **Sex** | **Primary vaccination** | **Third dose vaccination** | **Time of confirmed SARS-CoV-2 infection** |
| --- | --- | --- | --- | --- | --- |
| 19 | 26 | M | ChAdOx1 | BNT162b2 | 26 days before 6-months post-dose 3 sample |
| 20 | 27 | F | ChAdOx1 | BNT162b2 | 12 days before 1-month post-dose 3 sample |
| 21 | 29 | F | ChAdOx1 | BNT162b2 | 38 days before 6-months post-dose 3 sample |
| 22 | 32 | F | ChAdOx1 | BNT162b2 | NA |
| 23 | 33 | F | ChAdOx1 | BNT162b2 | NA |
| 24 | 33 | M | ChAdOx1 | BNT162b2 | Before 6-months post-dose 3 sample |
| 25 | 34 | F | ChAdOx1 | BNT162b2 | 67 days before 6-months post-dose 3 sample |
| 26 | 35 | M | ChAdOx1 | BNT162b2 | NA |
| 27 | 36 | F | ChAdOx1 | BNT162b2 | 42 days before 6-months post-dose 3 sample |
| 28 | 37 | F | ChAdOx1 | BNT162b2 | NA |
| 29 | 37 | F | ChAdOx1 | BNT162b2 | 20 days before 6-months post-dose 3 sample |
| 30 | 38 | F | ChAdOx1 | BNT162b2 | NA |
| 31 | 42 | M | ChAdOx1 | BNT162b2 | 61 days before 6-months post-dose 3 sample |
| 32 | 43 | F | ChAdOx1 | BNT162b2 | NA |
| 33 | 45 | F | ChAdOx1 | BNT162b2 | NA |
| 34 | 47 | F | ChAdOx1 | BNT162b2 | 10 days before 6-months post-dose 3 sample |
| 35 | 48 | F | ChAdOx1 | BNT162b2 | NA |
| 36 | 48 | F | ChAdOx1 | BNT162b2 | NA |
| 37 | 48 | F | ChAdOx1 | mRNA1273 | NA |
| 38 | 50 | F | ChAdOx1 | BNT162b2 | Infection confirmed, no record of date (between 1- and 6- months post dose 3) |
| 39 | 54 | F | ChAdOx1 | BNT162b2 | NA |
| 40 | 56 | F | ChAdOx1 | BNT162b2 | NA |
| 41 | 57 | M | ChAdOx1 | BNT162b2 | NA |
| 42 | 58 | F | ChAdOx1 | BNT162b2 | 15 days before 1-month post-dose 3 sample |
| 43 | 64 | F | ChAdOx1 | BNT162b2 | NA |
|  | 42 (26-64)  median w range | 80% Female |  | 96% BNT162b2 |  |

**Supplementary Table 3. Participant characteristics**

|  | **Homologous** | **Heterologous** | ***p*-value**^1^ |
| --- | --- | --- | --- |
| Timing of blood sampling (days post-vaccination; median w range) | | | |
| 3-4 weeks post-dose 1 | 23 (19-31) | 28 (25-29) | **0.0012** |
| 1-month post-dose 2 | 28 (25-35) | 28 (27-36) | 0.36 |
| 6-months post-dose 2 | 185 (181-194) | 178 (154-197) | **0.0003** |
| 1-month post-dose 3 | 31 (27-43) | 29 (28-64) | 0.69 |
| 6-months post-dose 3 | 184 (167-212) | 184 (129-208) | 0.92 |
| Age (years; median w range) | 37 (24-62) | 42 (26-64) | 0.15 |
| % Female | 55.6% (10/18) | 80% (20/25) | 0.09^2^ |
| BNT162b2 at dose 3 | 100% (18/18) | 96% (24/25) | 0.39^2^ |
| ^1^ Non-parametric Mann-Whitney test with Bonferroni correction for multiple comparisons  ^2^ Chi-square test | | | |

**Supplementary Table 4. Composition of the antibody panels**

|  | **Fluorochrome** | | | | | | | | | | | | | | | |
| --- | --- | --- | --- | --- | --- | --- | --- | --- | --- | --- | --- | --- | --- | --- | --- | --- |
| Tube | **BUV395** | **BUV496** | **BUV737** | **BV421** | **BV480** | **BV650** | **BV711** | **BV786** | **FITC** | **PerCP-Cy5.5/ BB700** | **PE** | **PE-Vio615** | **PC7/**  **PE-Cy7** | **APC** | **AF700** | **APC-H7** |
| 1. TruCount | **-** | **-** | **-** | **-** | **-** | **-** | **-** | - | CD3 | CD45 | CD16 + CD56 | **-** | CD4 | CD19 | - | CD8A |
| 2. Ag-specific Bmem | RBD  ancestral | CD3 | RBD ancestral | CD27 | RBD BA.2 | RBD BA.5 | CD21 | CD71 | IgG2 + IgG3 | IgD | IgG1 + IgG2 | IgA | CD19 | IgG4 | Viability | CD38 |
| 3. Streptavidin control | Strep | - | Strep | CD27 | Strep | Strep | - | - | CD3 | IgD | - | - | CD19 | - | Viability | - |

**Supplementary Table 5. Antibody list**

| **Marker** | **Fluorochrome** | **Clone** | **Supplier** | **Cat. number** | **Volume/**  **test (μl)** | **Tube(s)** |
| --- | --- | --- | --- | --- | --- | --- |
| CD3 | BUV496 | UCHT1 | BD Bioscience | 612940 | 1 | 2 |
| CD3 | FITC | SK7 | BD Biosciences | 555332 | 46 ng* | 1 |
| CD3 | FITC | UCHT1 | BD Biosciences | 662995 | 1 | 3 |
| CD4 | PE-Cy7 | SK3 | BD Biosciences | 555332 | 30 ng* | 1 |
| CD8A | APC-Cy7 | SK1 | BD Biosciences | 555332 | 126 ng* | 1 |
| CD16 | PE | B73.1 | BD Biosciences | 555332 | 33 ng* | 1 |
| CD19 | APC | SJ25C1 | BD Biosciences | 555332 | 46 ng* | 1 |
| CD19 | PE-Cy7 | SJ25C1 | BD Biosciences | 557835 | 5 | 2 |
| CD21 | BV711 | B-ly4 | BD Biosciences | 563163 | 5 | 2 |
| CD27 | BV421 | M-T271 | BD Biosciences | 562513 | 1 | 2 / 3 |
| CD38 | APC-H7 | HB7 | BD Biosciences | 303534 | 1 | 2 |
| CD45 | PerCP-Cy5.5 | 2D1 | BD Biosciences | 555332 | 120 ng* | 1 |
| CD56 | PE | NCAM16.2 | BD Biosciences | 555332 | 22 ng* | 1 |
| CD71 | BV786 | M-A712 | BD Biosciences | 563768 | 1 | 2 |
| IgA | PE-Vio615 | REA1014 | Miltenyi Biotec | 130-116-882 | 1.5 | 2 |
| IgD | BB700 | IA6-2 | BD Biosciences | 566538 | 1 | 2 |
| IgG1 | PE | G17-1 | BD Biosciences | 624049 | 0.1 | 2 |
| IgG2 | PE | HP6002 | BD Biosciences | 624049 | 0.5 | 2 |
| IgG2 | FITC | HP6002 | BD Biosciences | 624045 | 1 | 2 |
| IgG3 | FITC | HP6047 | BD Biosciences | 624045 | 0.5 | 2 |
| IgG4 | APC | SAG4 | Cytognos | CYT-IGG4AP | 2 | 2 |
| Strep | BUV395 | - | BD Biosciences | 564176 | 0.67 | 3 |
| Strep | BUV737 | - | BD Biosciences | 564293 | 0.67 | 3 |
| Strep | BV480 | - | BD Biosciences | 564876 | 0.67 | 3 |
| Strep | BV650 | - | Biolegend | 405232 | 0.13 | 3 |
| Viability | AF700 | - | BD Biosciences | 564997 | 0.1 | 2 / 3 |
| * total amount of antibody in ng per test | | | | | | |

**Supplementary Table 6. Flow cytometer set-up**

| **LSRFortessa X-20** | | **LSRII** | | **FACSLyric** | | **Fluorochromes used in this study** |
| --- | --- | --- | --- | --- | --- | --- |
| **355 nm** | | **-** | | **-** | |  |
| 379/28 | No LP | **-** | - | - | - | BUV395 |
| 525/50 | 505 LP | **-** | - | - | - | BUV496 |
| 740/35 | 690 LP | **-** | - | - | - | BUV737 |
| **405 nm** | | **405 nm** | | **405 nm** | |  |
| 450/50 | No LP | 450/50 | No LP | 448/45 | 448/45 | BV421 |
| 525/50 | 505 LP | 525/50 | 505 LP | 528/45 | 500 LP | BV480 |
| - | - | 586/15 | 570 LP | - | - | - |
| 610/20 | 600 LP | 610/20 | 600 LP | 606/36 | 606/36 | - |
| 670/30 | 635 LP | 660/20 | 630 LP | - | - | BV650 |
| 710/50 | 685 LP | 710/50 | 685 LP | 715/50 | 715/50 | BV711 |
| 780/60 | 750 LP | 780/60 | 750 LP | 755 LP | 755 LP | BV786 |
| **488 nm** |  | **488 nm** |  | **488 nm** | |  |
| 488/10 | No LP | 488/10 | No LP | 488/15 | No LP | SSC |
| 530/30 | 505 LP | 530/30 | 505 LP | 527/32 | 507 LP | FITC |
| **-** | **-** | **-** | **-** | 586/42 | 560 LP | PE |
| 710/50 | 685 LP | 710/50 | 630 LP | 700/54 | 665 LP | PerCP-Cy5.5, BB700 |
| **-** | **-** | **-** | **-** | 783/56 | 752 LP | PE-Cy7 |
| **561 nm** |  | **561 nm** |  | **-** | |  |
| 586/15 | No LP | 582/15 | No LP | - | - | PE |
| 610/20 | 600 LP | 610/20 | 600 LP | - | - | PE-Vio615 |
| 675/50 | 635 LP | 685/35 | 635 LP | - | - | - |
| 780/60 | 750 LP | 780/60 | 750 LP | - | - | PE-Cy7, PC7 |
| **640 nm** |  | **640 nm** |  | **640 nm** | |  |
| 670/30 | No LP | 670/14 | No LP | 660/10 | 660/10 | APC |
| 730/45 | 690 LP | 730/45 | 690 LP | 720/30 | 705 LP | Fixable Viability Stain 700 |
| 780/60 | 750 LP | 780/60 | 750 LP | 783/56 | 752 LP | APC-H7, APC-Cy7 |

**Supplementary Table 7. Target values for 7^th^ peak of rainbow beads in fluorescent channels**

| **Fluorochrome** | **Channel** | Lower (-15%) | **Target MFI** | Upper (+15%) | Recommendation |
| --- | --- | --- | --- | --- | --- |
| **BUV395** | **UV395** | 17,000 | **20,000** | 23,000 | In-house |
| **BUV496** | **UV525** | 23,800 | **28,000** | 32,200 | In-house |
| **BUV737** | **UV737** | 21,2500 | **25,000** | 28,750 | In-house |
| **BV421** | **V450** | 100,452 | **118,178** | 135,905 | EuroFlow |
| **BV480** | **V525** | 93,871 | **110,436** | 127,002 | EuroFlow |
| **BV650** | **V670** | 34,383 | **40,450** | 46,518 | In-house |
| **BV711** | **V710** | 15,079 | **17,740** | 20,401 | In-house |
| **BV786** | **V780** | 1,903 | **2,239** | 2,575 | In-house |
| **FITC** | **B530** | 28,752 | **33,826** | 38,900 | EuroFlow |
| **PerCP-Cy5.5/ BB700** | **B710** | 66,846 | **78,642** | 90,438 | EuroFlow |
| **PE** | **YG586** | 32,381 | **38,095** | 43,809 | EuroFlow |
| **PE-Vio615** | **YG610** | 178,500 | **210,000** | 241,500 | In-house |
| **PE-Cy7** | **YG780** | 8,316 | **9,783** | 11,250 | EuroFlow |
| **APC** | **R670** | 158,639 | **186,634** | 214,629 | EuroFlow |
| **AF700** | **R730** | 121,550 | **143,000** | 164,450 | In-house |
| **APC-H7** | **R780** | 64,194 | **75,522** | 86,850 | EuroFlow |
| Spherotech Rainbow Calibration particles (8 peaks) 3.41µm; cat nr. RCP-30-5A, Lot No. EAG01  As per EuroFlow recommendation (31). | | | | | |

**SUPPLEMENTARY FIGURES (n=8)**

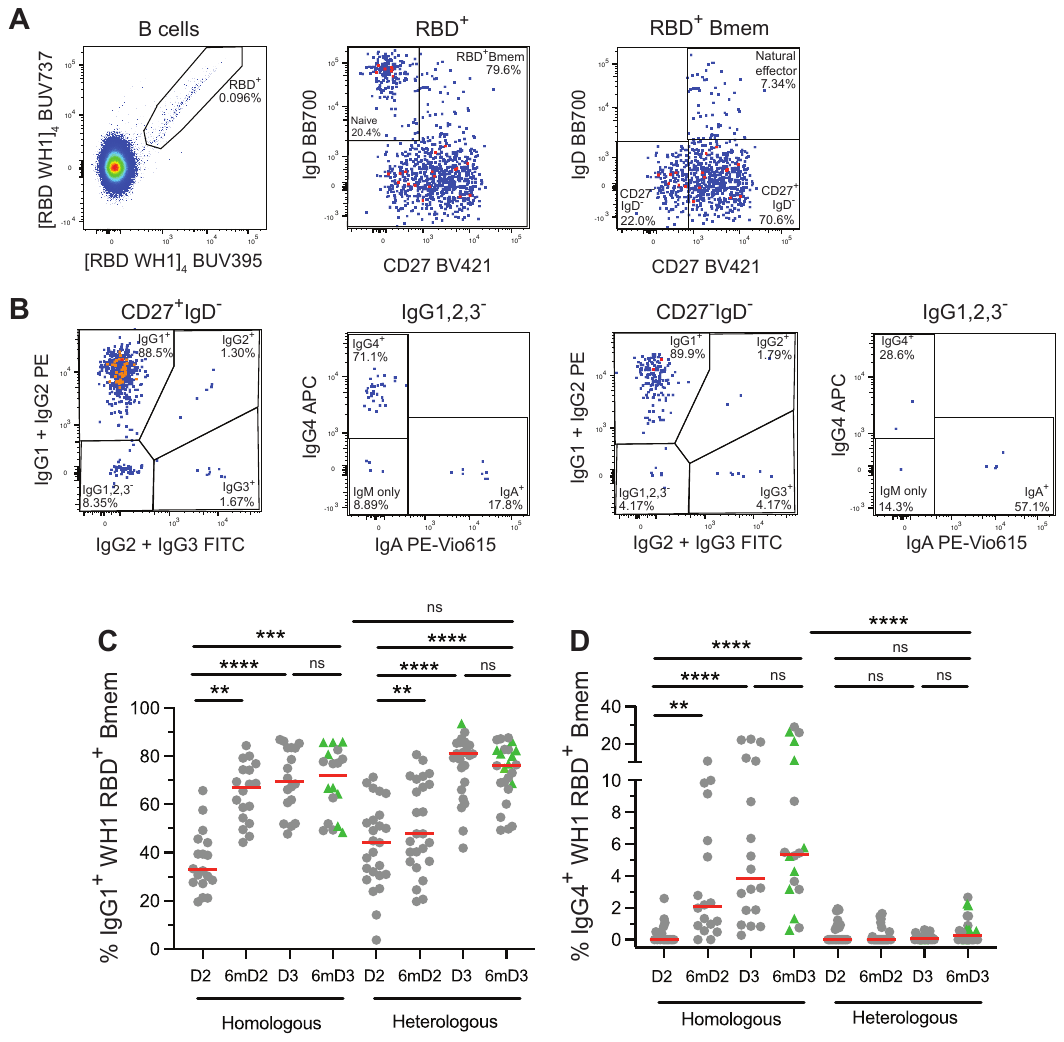

**Supplementary Figure 1: Predominant IgG1^+^ ancestral (WH1) RBD-specific Bmem response following homologous and heterologous vaccination.** (**A**) Identification of WH1 RBD-specific B cells, Bmem and CD27^+^ memory. (**B**) CD27^+/-^IgD^-^ RBD-specific Bmem were further separated based on IgG1,2,3,4 and IgA expression. Frequencies of (**C**) IgG1^+^ and (**D**) IgG4^+^ RBD-specific Bmem following homologous or heterologous vaccination. Green triangles represent individuals who had a confirmed breakthrough infection (BTI) prior to sampling (**Supplementary Tables 1 and 2**). Red lines in panels **C** and **D** represent median values. Kruskal-Wallis test with Dunn’s multiple comparisons test. ** *p* > 0.01, *** *p* > 0.001, **** *p* > 0.0001.

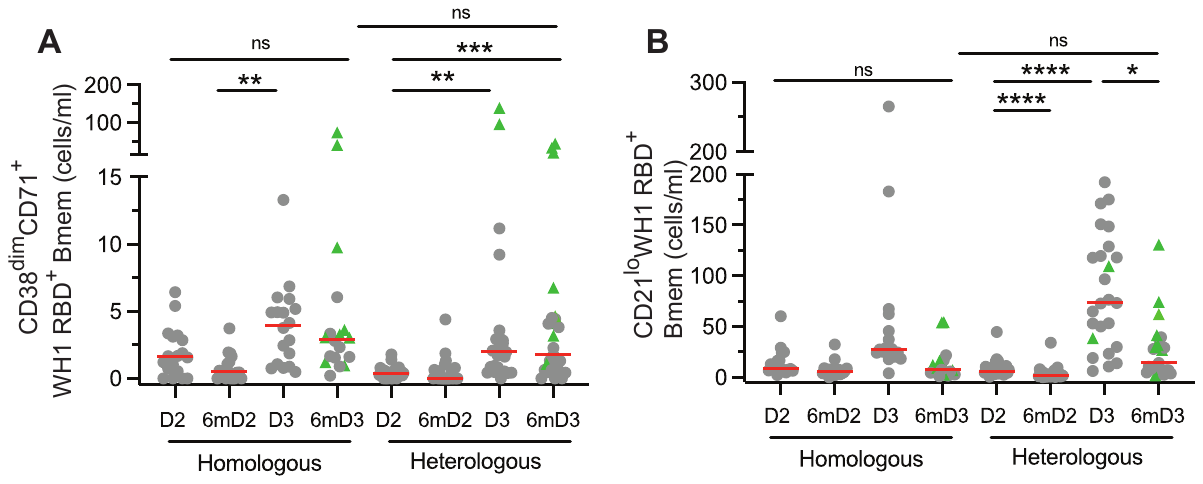

**Supplementary Figure 2: Extended phenotype of ancestral (WH1) RBD-specific Bmem following homologous or heterologous vaccination.** Absolute numbers of (**A**) CD38^dim^CD71^+^ and (**B**) CD21^lo^ RBD-specific Bmem. Green triangles represent individuals who had a confirmed breakthrough infection (BTI) prior to sampling (**Supplementary Tables 1 and 2**). Red lines represent median values. Kruskal-Wallis test with Dunn’s multiple comparisons test. * *p* > 0.05, ** *p* > 0.01, *** *p* > 0.001, **** *p* > 0.0001.

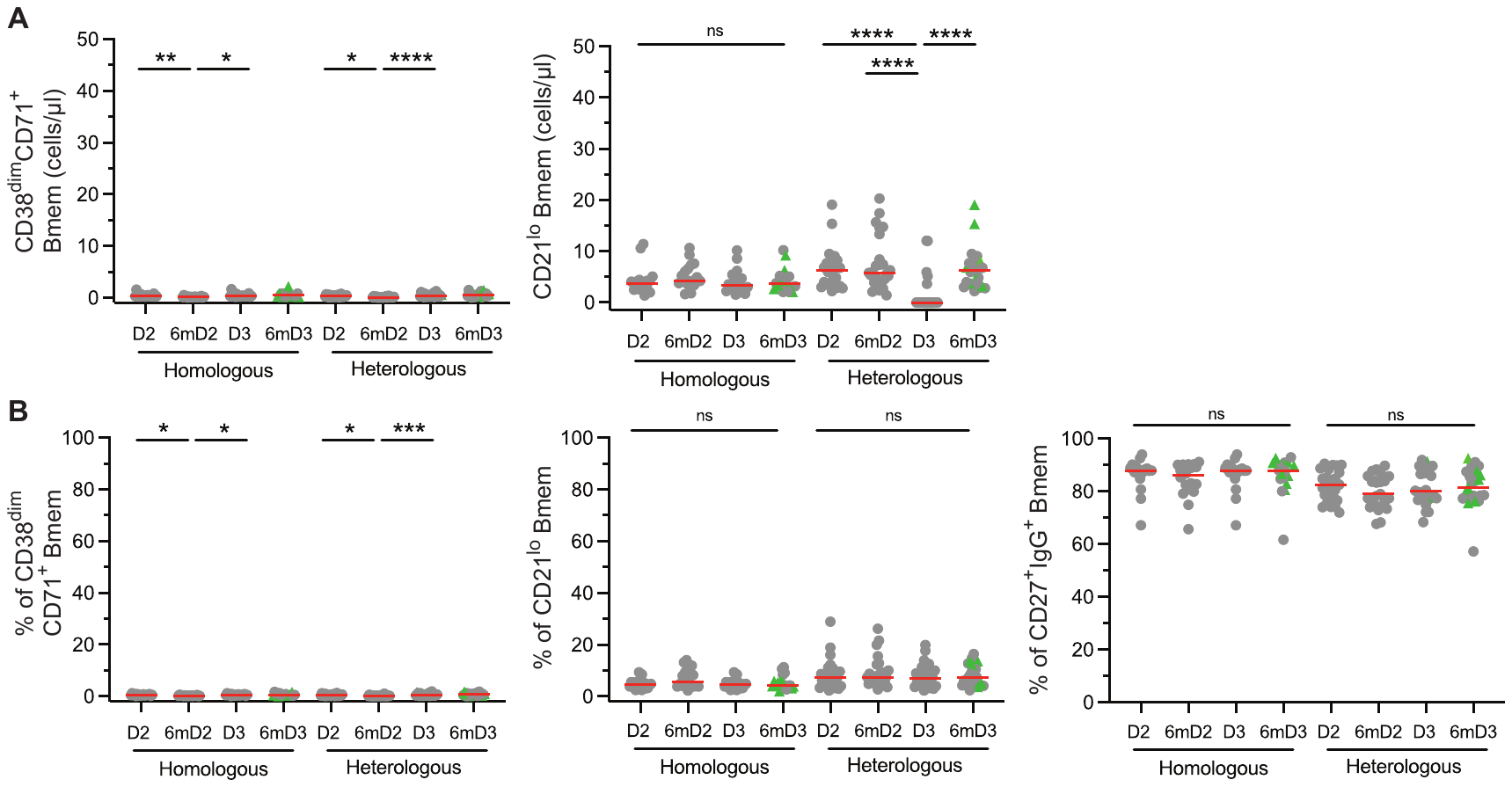

**Supplementary Figure 3: Extended phenotype of total Bmem following homologous or heterologous vaccination.** (**A**) Absolute numbers of CD38^dim^CD71^+^ and CD21^lo^ total Bmem Bmem. (**B**) Frequencies of CD38^dim^CD71^+^, CD21^lo^ and CD27^+^IgG^+^ total Bmem. Green triangles represent individuals who had a confirmed breakthrough infection (BTI) prior to sampling (**Supplementary Tables 1 and 2**). Red lines represent median values. Kruskal-Wallis test with Dunn’s multiple comparisons test. * *p* > 0.05, ** *p* > 0.01, ** *p* > 0.001, **** *p* > 0.0001.

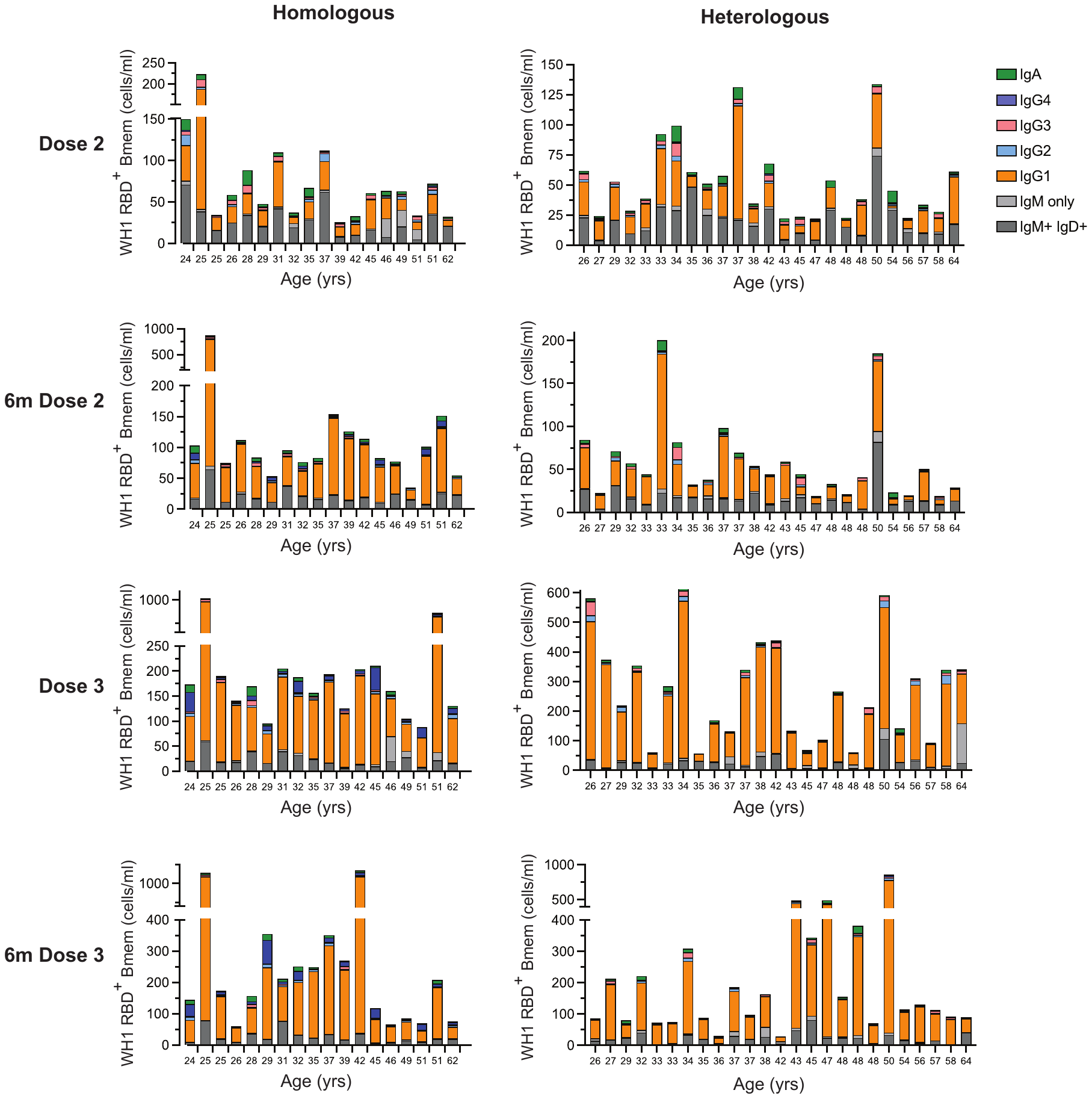

**Supplementary Figure 4: Individual ancestral (WH1) RBD-specific Bmem numbers following homologous or heterologous vaccination.** Absolute number of RBD-specific Bmem in each individual for the homologous and heterologous vaccination cohorts post-dose 2, 6-months post-dose 2, post-dose 3 and 6-months post-dose 3.

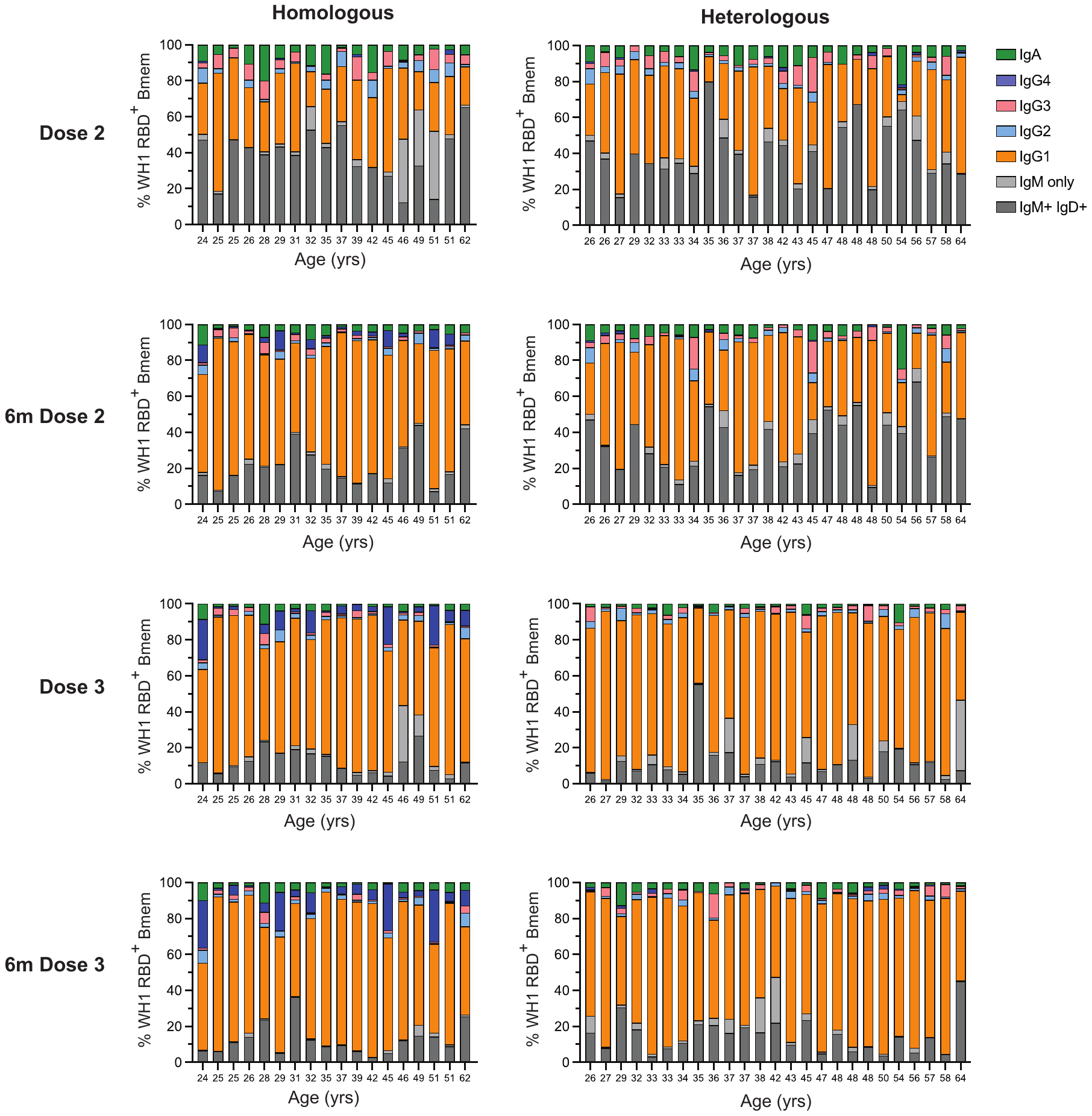

**Supplementary Figure 5: Individual frequencies of ancestral (WH1) RBD-specific Bmem following homologous or heterologous vaccination.** Frequencies of subsets within WH1 RBD-specific Bmem in each individual for the homologous and heterologous vaccination cohorts 1-month post-dose 2, 6-months post-dose 2, 1-month post-dose 3 and 6-months post-dose 3.

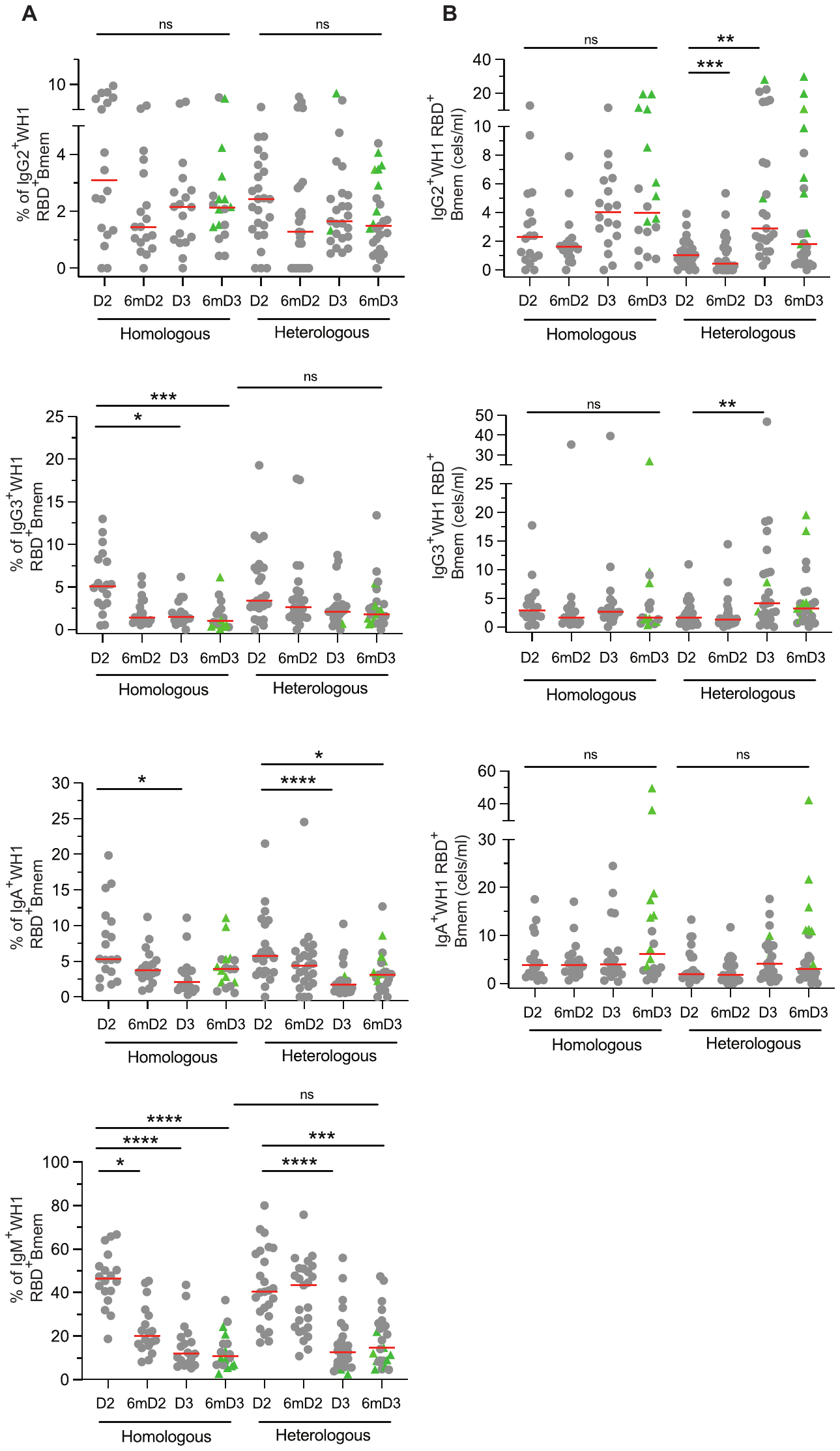

**Supplementary Figure 6: Ancestral (WH1) RBD-specific Bmem subsets following homologous or heterologous vaccination.** (**A**) Frequencies and (**B**) absolute numbers of IgG2^+^, IgG3^+^, IgA^+^ and IgM^+^ WH1 RBD-specific Bmem. Green triangles represent individuals who had a confirmed breakthrough infection (BTI) prior to sampling (**Supplementary Tables 1 and 2**). Red lines represent median values. Kruskal-Wallis test with Dunn’s multiple comparisons test. * *p* > 0.05, ** *p* > 0.01, ** *p* > 0.001, **** *p* > 0.0001.

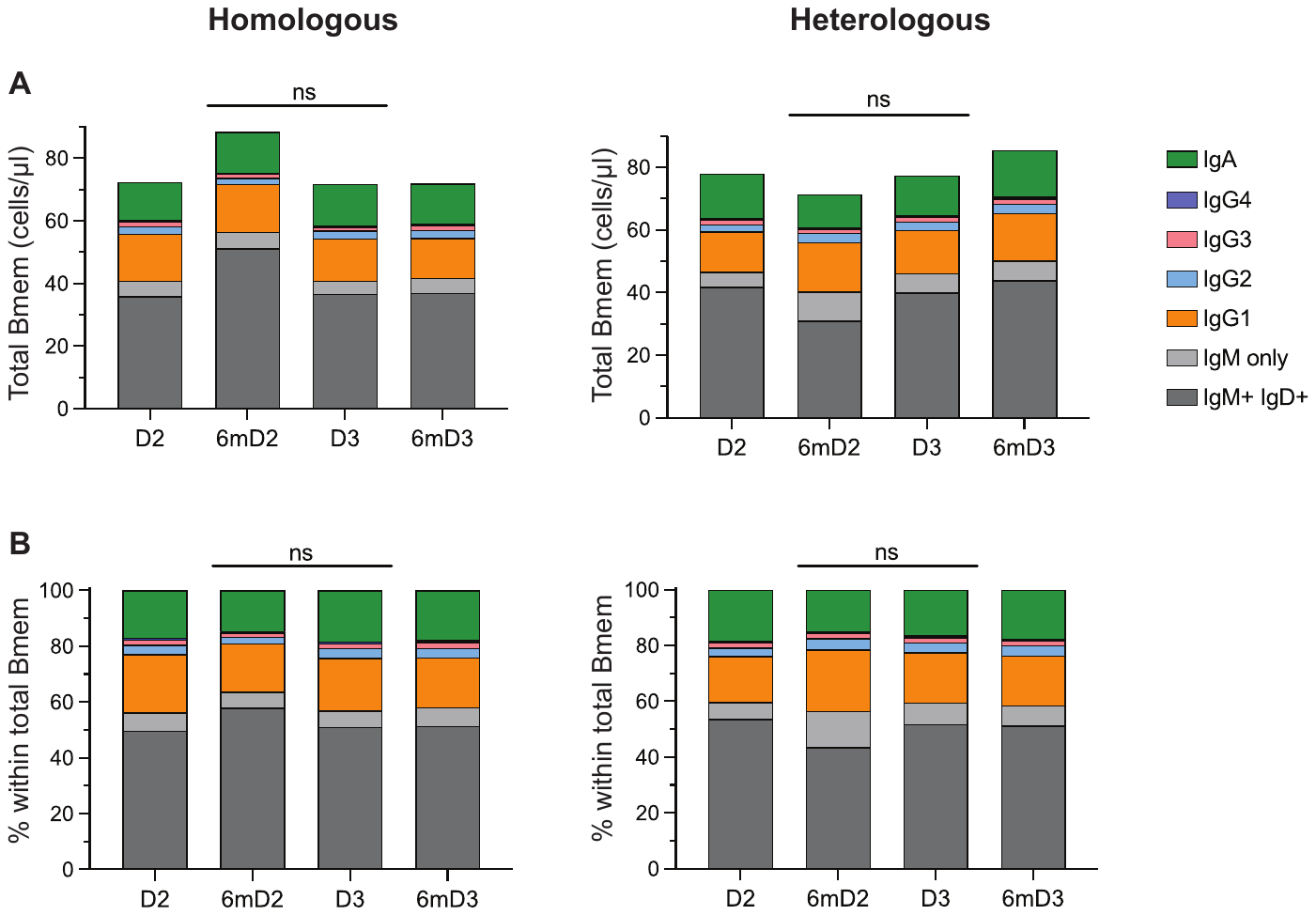

**Supplementary Figure 7: Absolute numbers and frequencies of total Bmem Ig isotypes and IgG subclasses following homologous or heterologous vaccination.** Median values of (**A**) absolute numbers and (**B**) frequencies of total Bmem following homologous or heterologous vaccination. Kruskal-Wallis test with Dunn’s multiple comparisons test.

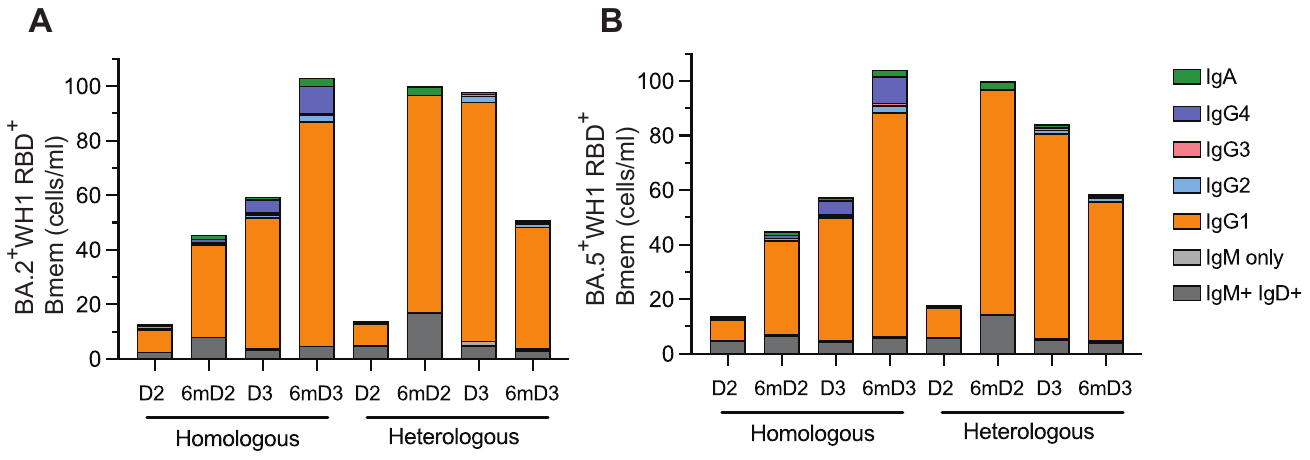

**Supplementary Figure 8: Immunophenotype of variant binding RBD-specific Bmem.** Median absolute numbers of ancestral (WH1) RBD-specific Bmem that also recognize Omicron (**A**) BA.2 or (**B**) BA.5.
